# The radish Ogura fertility restorer impedes translation elongation along its cognate CMS-causing mRNA

**DOI:** 10.1101/2021.03.17.435859

**Authors:** Chuande Wang, Lina Lezhneva, Nadège Arnal, Martine Quadrado, Hakim Mireau

## Abstract

The control of mRNA translation has been increasingly recognized as a key regulatory step for gene control but clear examples in eukaryotes are still scarce. Nucleo-cytoplasmic male sterilities (CMS) represent ideal genetic models to dissect genetic interactions between the mitochondria and the nucleus in plants. This trait is determined by specific mitochondrial genes and is associated with a pollen sterility phenotype that can be suppressed by nuclear genes known as restorer-of-fertility (*Rf*) genes. In the study, we focused on the Ogura CMS system in rapeseed and showed that the suppression to male sterility by the PPR-B fertility restorer (also called Rfo) occurs through a specific inhibition of the translation of the mitochondria-encoded CMS-causing mRNA *orf138*. We also demonstrate that PPR-B binds within the coding sequence of *orf138* and acts as a ribosome blocker to specifically impede translation elongation along the *orf138* mRNA. Rfo is the first recognized fertility restorer shown to act this way. These observations will certainly facilitate the development of synthetic fertility restorers for CMS systems in which efficient natural Rfs are lacking.

## Introduction

Fine-tuning of gene expression provides cells with necessary proteins to function properly. Each step in the flow of information going from DNA to proteins offers cells with potential checkpoints to adjust the type and the activity of proteins they synthesize. Changes in transcriptional patterns play major roles in gene regulation in both prokaryotes and eukaryotes and is orchestrated by different molecular means. Posttranscriptional regulatory mechanisms allow faster reshaping of cellular proteomes compared to purely transcriptional events. In particular, the control of mRNA translation has been increasingly recognized as a key regulatory step of gene control, in most genetic systems. All phases of translation, including initiation, elongation, termination and ribosome recycling constitute potential checkpoints to modulate gene expression. Translational control can be mediated by mRNA structural features or through the action of proteinaceous or RNA *trans-factors* (1–4).

In eukaryotic cells, the spread-out of genetic information between the nuclear and cytoplasmic genomes adds an additional layer of complexity to gene regulation processes. Cytoplasmic genomes are extremely low in gene contents and virtually all regulatory functions of organellar genes expression are nuclear encoded (5–7). Nucleo-cytoplasmic male sterilities (CMS) represent ideal genetic models to understand nucleo-mitochondrial coadaptation processes. CMS is a widely expanded trait of plants characterized by an inability of plant to produce functional pollen. CMS traits are specified by poorly-conserved mitochondrial genes and can be suppressed by nuclear-encoded restorer of fertility (*Rf*) genes that specifically act to in most cases down-regulate the expression of corresponding CMS-specifying mitochondrial genes (8). In the last years, several *Rf* genes were identified in various crop species and most of them were found to encode proteins belonging to the large family of pentatrico-peptide repeat (PPR) proteins (9). PPR proteins are highly specific RNA-binding proteins that widely diversified in eukaryotes, mainly in terrestrial plants (10). PPR proteins have been shown to play multifarious roles in mitochondrial and plastid RNA expression processes, going from gene transcription to mRNA translation (11). Rf-PPR mediated suppressing activity often alters mitochondrial CMS-causing mRNA levels (12–15). The Ogura CMS Rf-PPR from radish stands as an exception among fertility restores as it was shown to not affect its cognate CMS-conferring mRNA, either in size or in abundance (16). The Ogura CMS, originally identified in radish (*Raphanus sativus*) and later transferred to rapeseed (*Brassica napus),* is controlled by the mitochondrial *orf138* locus (17, 18). We showed that the Ogura restorer of fertility protein named PPR-B (19–21) associates *in vivo* with the *orf138* mRNA and that this association leads to a strong decrease in the Orf138 protein level, notably in tapetal cells and developing pollen grains (22). The Ogura Rf-PPR was thus suspected to impact the translation of *orf138* mRNA, but this hypothesis needed to be validated and the way by which PPR-B may interfere with the translation of the *orf138* transcript determined. In this present study, we demonstrate that PPR-B CMS-suppressing activity implies a specific down-regulation of *orf138* translation and, very interestingly, that this control operates through a blockade of ribosome progression along the *orf138* coding sequence. The Ogura fertility restorer is the first Rf protein demonstrated to act this way.

## Results

### The Orf138 protein is not produced in mitochondria in the presence of PPR-B

The biochemical characterization of fertility restorers *in planta* has often been rendered difficult by the tissue-specificity of associated molecular mechanisms (23). This is true for the Ogura CMS system, as we showed that fertility restoration correlates with a profound decrease in Orf138 in tapetal cells and microspores but not in other plant tissues (22). We could, however, produce a restored rapeseed transgenic line (named B1) containing four copies of *PPR-B* and in which Orf138 reduction was nearly ubiquitous. This provided us with an ideal biological material to characterize the effect of PPR-B on Orf138 production since the observed decrease was homogeneous across all plant tissues in this line. Immunoblot analyses were first conducted to confirm the near-complete disappearance of Orf138 in mitochondrial extracts prepared from B1 plant inflorescences (Figure 1A). *In-organello* protein syntheses in the presence of [^35^S] methionine were then carried out with mitochondria prepared from male-sterile (CMS) and restored B1 plants. Virtually identical translation profiles were revealed with both lines, except for one protein, close to 20 kDa, which was clearly visible in the CMS but not in B1 line (Figure 1B). A CMS-specific protein of the exact same size was identified in previous *in organello* profiles and was demonstrated to correspond to the Orf138 protein as it could be immunoprecipitated with an Orf138-specific antibody (24). The lack of Orf138 in B1 *in organello* translation products strongly favoured an incapacity of mitochondria to produce the Orf138 protein under the action of PPR-B, rather than an increased instability of Orf138 in the restoration context.

**Figure 1.**
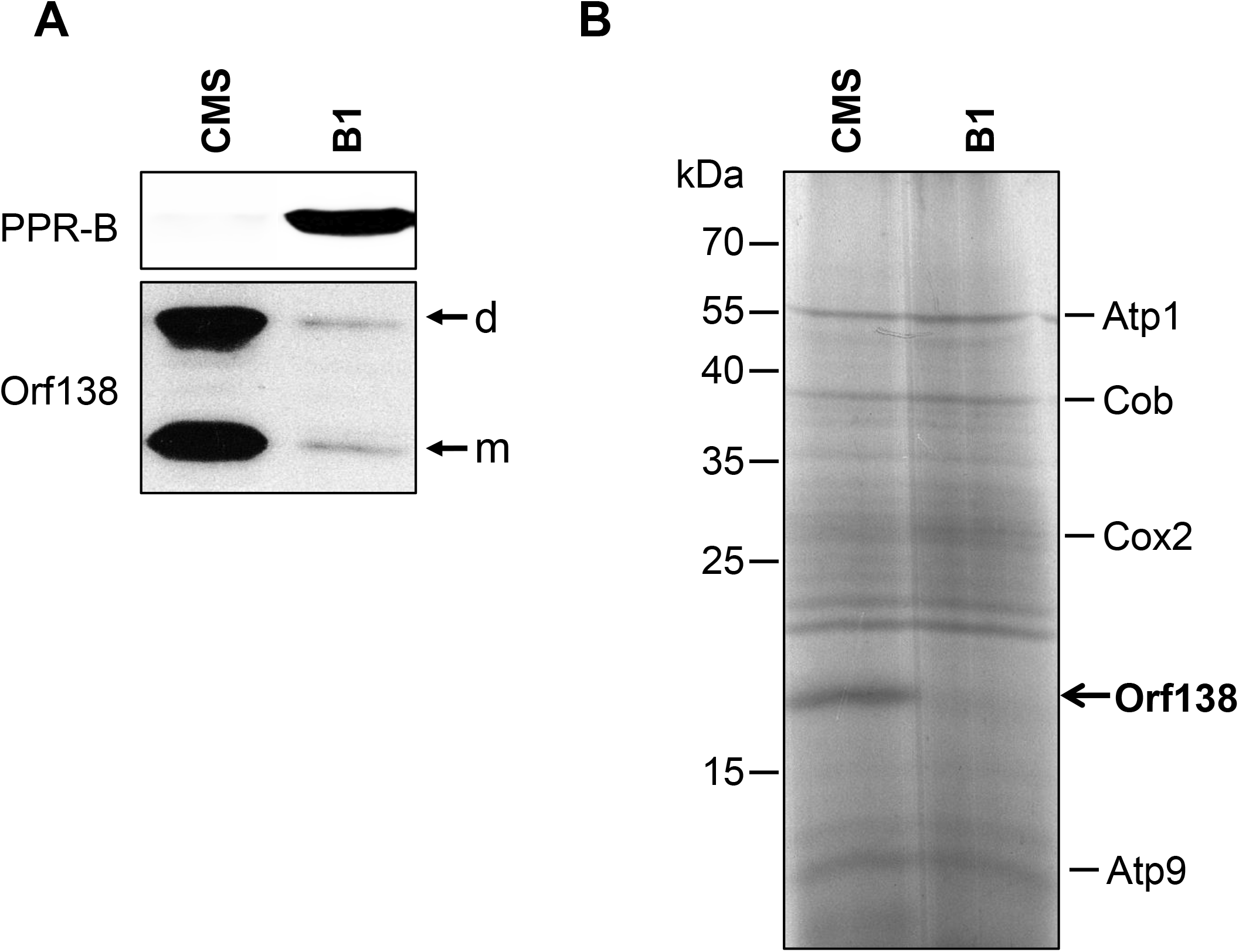
The Orf138 protein is not produced in the presence of PPR-B. (A) Immunoblot assay showing the steady-state levels of Orf138 and PPR-B proteins in floral tissues of male sterile (CMS) and transgenic B1 lines. m: monomeric Orf138, d: dimeric Orf138. (B) Autoradiograph of *in organello* [^35^S]methionine-labelled translation products from mitochondria isolated from male sterile (CMS) and transgenic B1 lines. Mitochondria translation products were separated on a 15% SDS-PAGE gel. The position of the Orf138 protein is indicated.

### The translation of *orf138* mRNA is impaired in the presence of PPR-B

The *in-organello* translation results strongly suggested a likely negative impact of PPR-B on the translational capacity of the *orf138* mRNA. Therefore, the translation status of the *orf138* transcript was first evaluated by polysome sedimentation analysis in both CMS and restored plants. The co-transcription of *orf138* with the *atp8* gene necessitated first to replace the cytoplasm of the B1 line with that of the male-sterile cybrid 18S line in which the *orf138* gene is not associated with *atp8* and transcribed as a monocistronic mRNA (24). The B1 line was then used to pollinate male-sterile 18S flowers and several F1 descendants were tested by immunoblot analysis to evaluate their content in Orf138 protein accumulation. In control, a non-transgenic Pactol Fertile (PF) X 18S F1 hybrid line was also generated. Unlike male-sterile PF/18S plants, all B1/18S hybrid plants were found to accumulate barely-detectable levels of Orf138, as in the original B1 line (*SI Appendix,* Fig. S1). Polysomes were then isolated from rapeseed B1/18S and PF/18S inflorescences and fractionated on continuous sucrose density gradients. Ten fractions were collected along the gradients after centrifugation and analysed by subsequent RNA gel blot. Polysome integrity was verified by the distribution of ribosomal RNAs along the gradients in the presence of MgCl2 (*SI Appendix*, Fig. S2). The disruption of polysomes with EDTA indicated that polysomal RNAs migrated toward the centre and the bottom of the gradients, whereas ribosome-free mRNAs accumulated in the upper fractions (*SI Appendix*, Fig. S2). Total RNAs were extracted from each fraction and subjected to RNA gel-blot analysis using probes specifically recognizing the *orf138* transcript, as well as *atp9* and *atp1* as controls (Fig. 2*A*). Obtained hybridization signals were then quantified and their relative distribution along the gradients was determined for each transcript (Fig. 2*B*). In the absence of PPR-B, we observed that the majority of the *orf138* signal accumulated in the central fractions, suggesting active translation of *orf138* in this genetic context. However in the presence of PPR-B, the pick of *orf138* hybridization signal was clearly shifted toward the upper fractions revealing a negative impact of PPR-B on *orf138* and polysome association. The distributions of *atp1* and *atp9* were unaffected by the PPR-B status, indicating that the PPR-B mediated translational impairment was specific to *orf138.* In next effort to better understand the origin of PPR-B translational repression, Ribo-Seq analyses were developed to compare the translational status of all mitochondria-encoded mRNA in CMS and fertility-restored lines. Total ribosome footprints were prepared from both lines and then mapped to the rapeseed mitochondrial genome and the *orf138* locus. RNA-Seq experiments were also developed to quantify the steady-state levels of all mitochondrial transcripts in the two genetic backgrounds. RNA-Seq data revealed no major impact of PPR-B on most mitochondrial mRNA abundance, except for several ribosomal protein transcripts whose steady-state levels were reduced by a factor of around 2 (*SI Appendix,* Fig. S3). Calculated translational efficiencies (see material and method for details) indicated a slight decrease of ribosome coverage for most mitochondrial transcript by less than a factor 2 in the restored B1 line compared to the CMS line, except for the *orf138* mRNAs which was found to around 16-fold less translated under PPR-B action (Fig. 3*A*). Interestingly, a few ribosomal protein transcripts (*e.g. rpl16* or *rps14*) appeared to be slightly up-translated in the presence of PPR-B. The impact of measured translational differences on mitochondrial protein accumulation was next evaluated by immunoblot assays (Fig. 3*B*). Among the few tested proteins, no major differences in protein steady-sate levels could be detected between CMS and B1 lines. The only reproducible differences concerned the Nad7 protein which appeared to slightly over-accumulate in B1 plants compared to the CMS line and, of course, the *orf138* protein which was here again hardly detectable in the B1 line. Altogether, these results strongly supported that the lack of Orf138 production in the restored B1 line resulted from a potent impairment of *orf138* mRNA translation.

**Figure 2.**
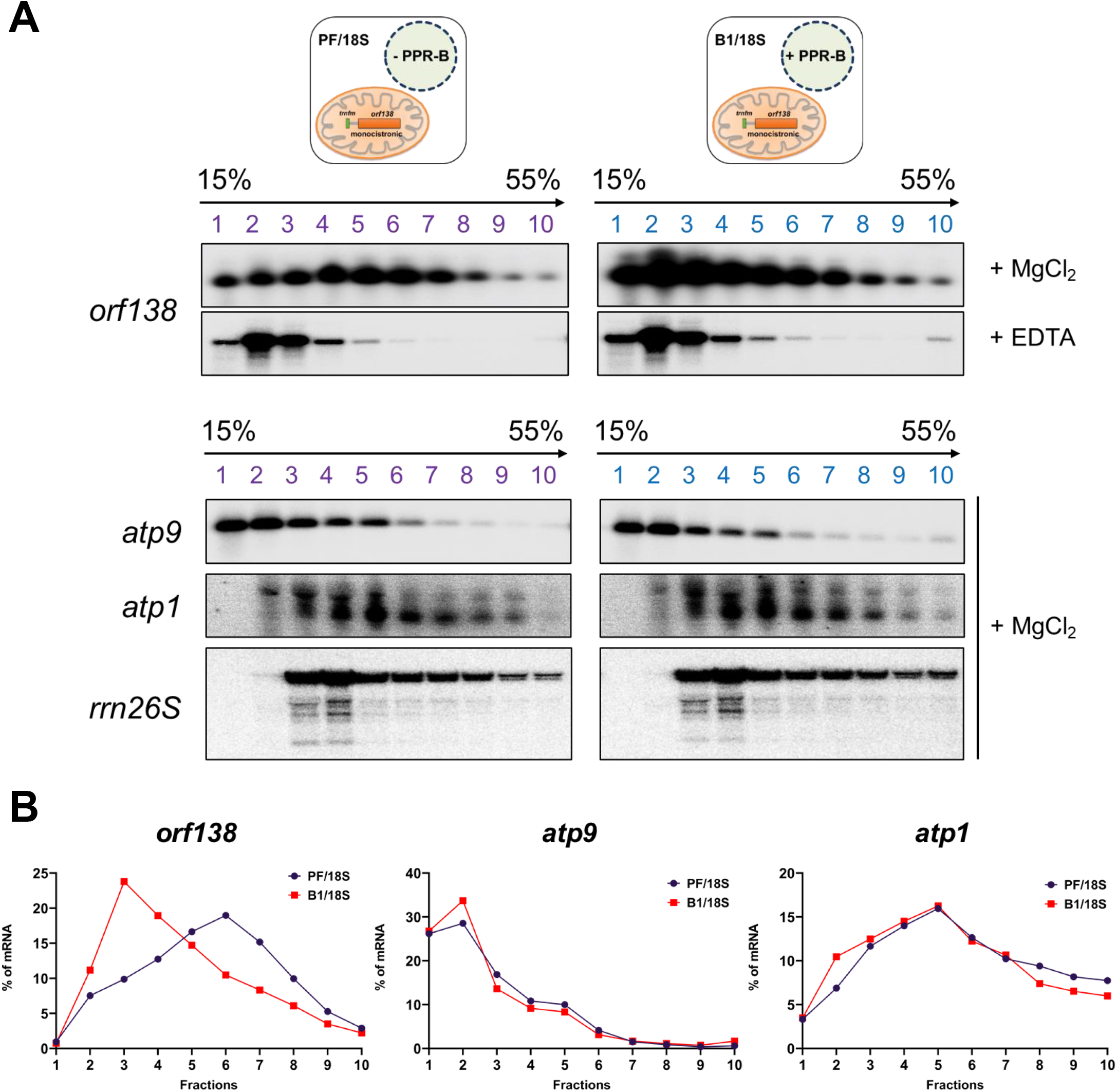
The association of the *orf138* transcript with mitochondrial polysomes is perturbed by PPR-B. (A) Total polysomes extracted from PF/18S and B1/18S hybrid plants were fractionated in 15% to 55% sucrose density gradients by ultracentrifugation and under conditions maintaining (+ MgCl2) or disrupting (+ EDTA) polysome integrity and analysed by RNA-gel blot assays using the indicated gene probes. rrn26S corresponds to the mitochondrial 26S ribosomal RNA and its hybridization profile attests for the integrity of polysomes along the gradients in the presence of MgCl_2_. (B) Quantification of hybridization signals along the polysomal gradients. The hybridization signals corresponding to *orf138*, *atp9* and *atp1* transcripts were quantified using the ImageQuant software (GE Healthcare Life Sciences) in each fraction and genotype. The indicated values correspond to the percentage of contribution of each fraction to the sum of all hybridization signals obtained over the entire gradients.

**Figure 3.**
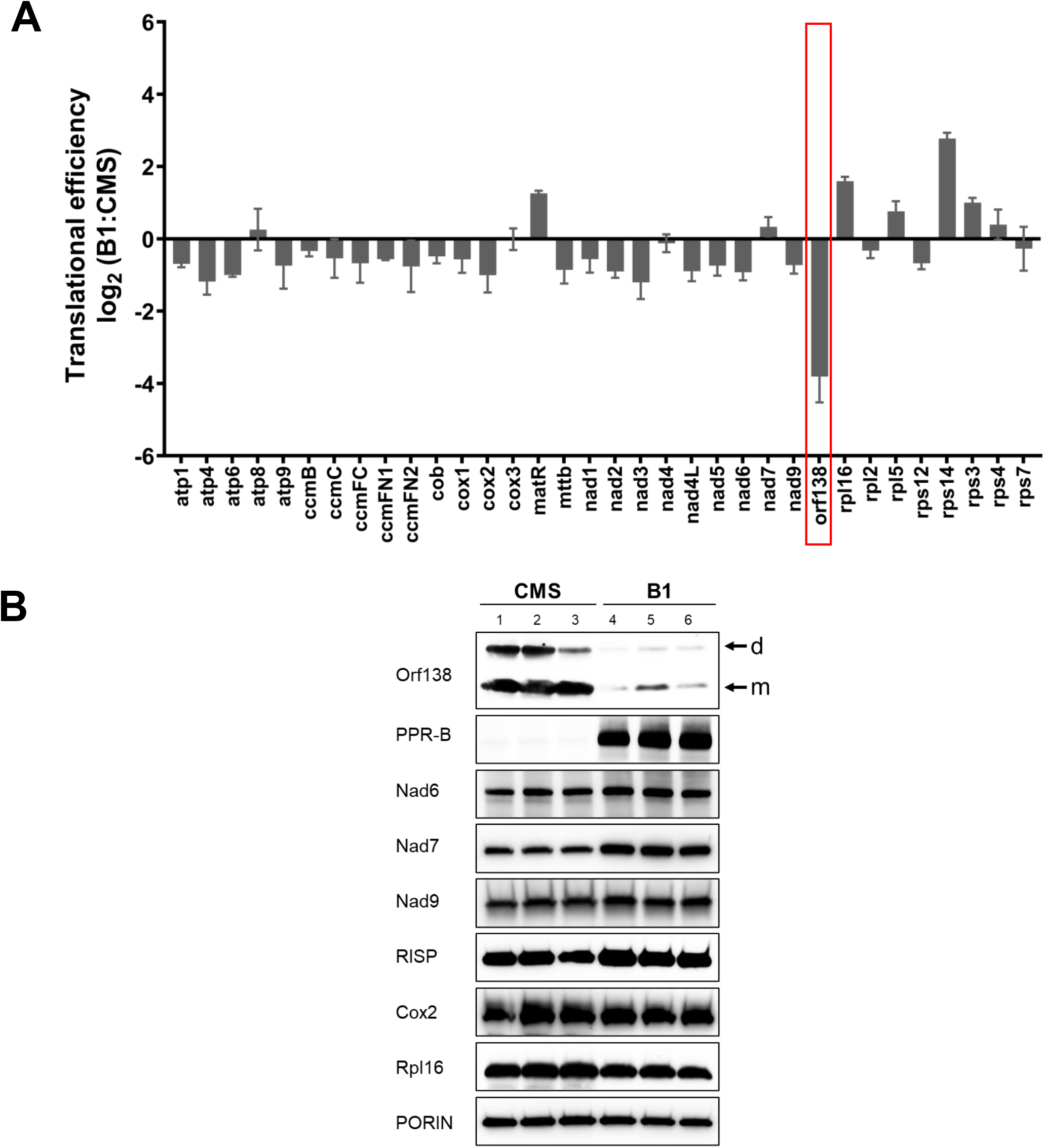
Translation of the *orf138* mRNA is strongly reduced under the effect of PPR-B. (A) Compared Ribo-Seq analysis of mitochondrial mRNAs in B1 and CMS lines. The bars depict log2 ratios of translational efficiencies of mitochondria-encoded mRNAs in B1 plants to the CMS line (see Material and Methods for details). The reported values are means of two independent biological replicates (error bars indicate SD). (B) SDS/PAGE immunoblots performed on total mitochondrial protein extracts prepared from flower buds of B1 and CMS plants and probed with antibodies to Orf138 and PPR-B as well as subunits of respiratory complex I (Nad6, Nad7 and Nad9), complex III (RISP), complex IV (Cox2) and the ribosomal protein Rpl16. Porin was used as protein loading control. Results obtained on three independent protein preparations are shown for each line. m: monomeric Orf138, d: dimeric Orf138.

### The PPR-B fertility restorer binds within the coding sequence of the *orf138* mRNA

We previously demonstrated that PPR-B specifically associates with the *orf138* mRNA *in vivo* (22). To understand how this association could negatively impact *orf138* translation, we sought to identify the binding site of PPR-B within the *orf138* transcript. PPR proteins are known to associate with their RNA target *via* a one PPR motif – one nucleotide recognition rule and that amino acid combinations at positions 5 and 35 of each repeat are major determinants for RNA base selection (25–27). We thus used the established PPR recognition code to predict the most likely binding regions of PPR-B within the rapeseed mitochondrial genome and the *orf138* locus (Fig. 4). Interestingly, a highly-probable binding region corresponding to a short segment located 55 nucleotides downstream of *orf138* AUG codon could be identified (Fig. 4*B*). Other identified potential binding sites located in intergenic regions of the *B. napus* mitochondrial genome and had virtually no chance to be associated with PPR-B-mediated restoration activity. To see if PPR-B showed significant affinity for the potential target identified in *orf138*, gel shift assays were developed using a series of *in-vitro* transcribed probes mapping to the 5’ region of the *orf138* mRNA and a recombinant form of PPR-B fused to the maltose binding protein (Fig. 5*A*). Among the different RNA probes tested, PPR-B showed a clear and strong binding affinity for all the probes containing the predicted binding site (Figs. 5*B* and *5C*). In contrast, PPR-B did not associate with nearby probes that did not contain the GTAAAGTTAGTGTAATA sequence, strongly supporting the binding predictions and that this sequence represented the PPR-B binding site within the *orf138* transcript.

**Figure 4.**
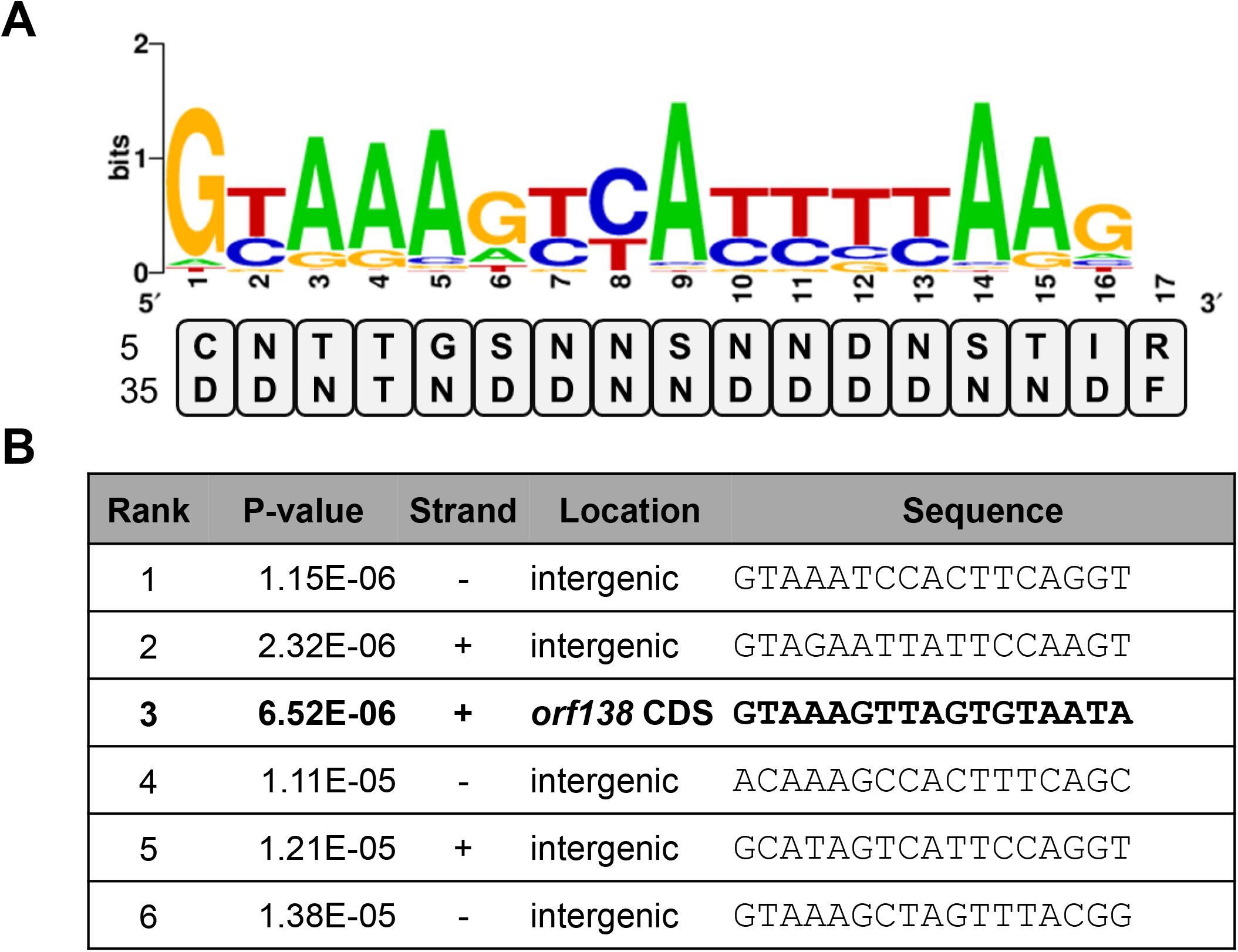
The PPR-B protein is predicted to associate within the coding sequence of *orf138*. (A) Combinations of amino acids at position 5 and 35 of each PPR-B PPR repeat are listed from N to C-terminus. The generated combinations were used to calculate probabilities of RNA base recognition by each PPR-B PPR repeat according to the PPR code (25–27). The sequence logo depicting these probabilities was obtained with http://weblogo.berkeley.edu/. (B) The nucleotide preference scores were then used to scan both strands of the *B. napus* mitochondrial genome (GenBank AP006444.1) and the sequence of the *orf138* locus (18) with the FIMO program. The sequence of the six most-probable PPR-B binding sequences are shown with their respective location. The *P*-values were determined with the FIMO program.

**Figure 5.**
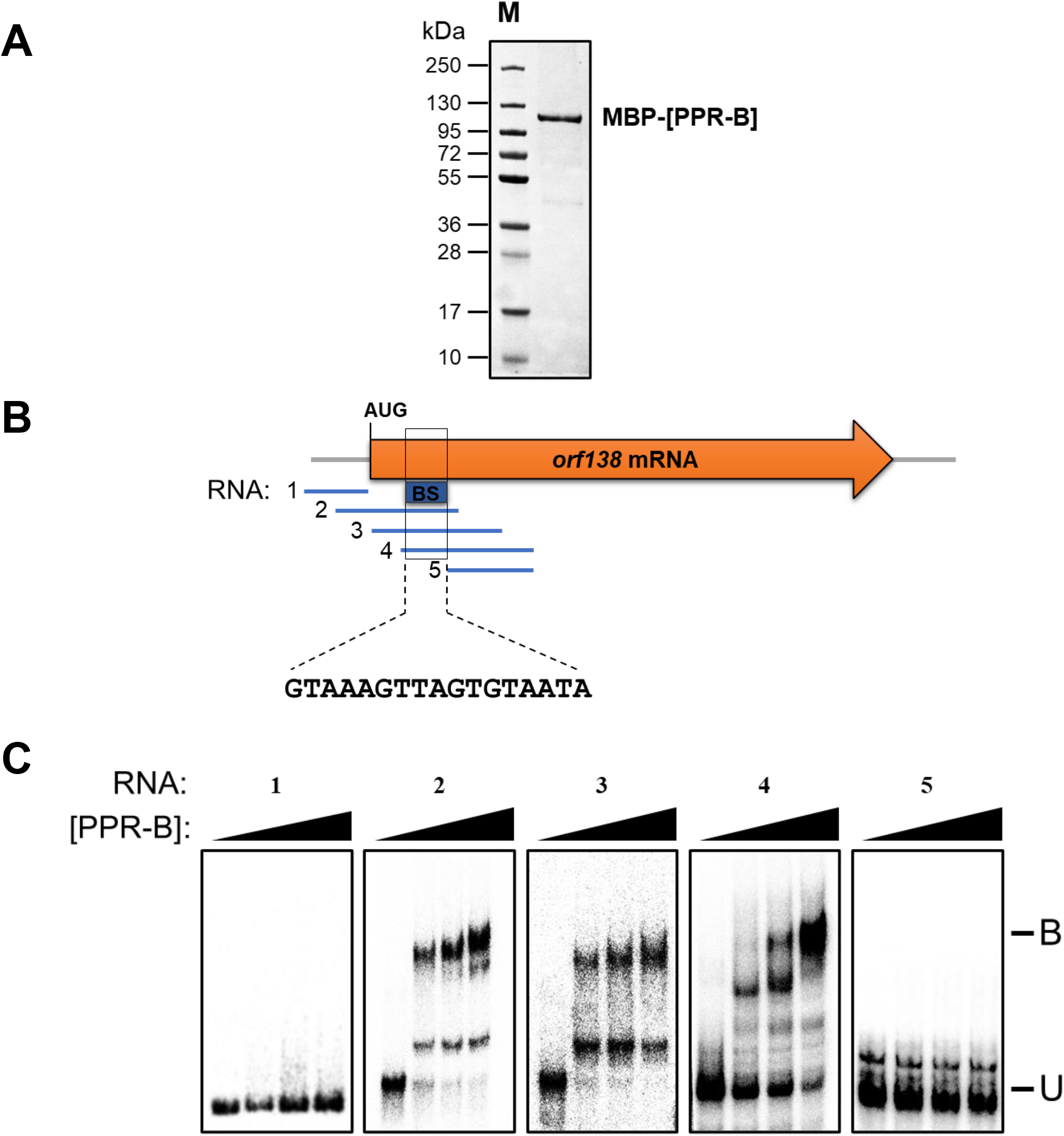
PPR-B binding binds 55 nucleotides downstream of the *orf138* translational start codon. (A) SDS-PAGE gel stained with Coomassie blue demonstrating the purity of the MBP-PPR-B fusion expressed and purified from *E. coli.* Five micrograms of purified MBP-PPR-B were loaded on the shown gel. Protein molecular weight markers are shown (M). (B) Schematic repression of the relative positions of the different *in-vitro* transcribed RNA probes used in gel shift assays shown in panel C. The location and sequence of the predicted PPR-B binding site (BS) are also displayed. (C) Gel mobility shift assays done with the MBP-PPR-B fusion and the RNA probes shown in panel B. 200 pM of radiolabeled RNA probes along with 0 to 800 nM of the fusion protein and 0.5 mg/ml heparin as negative binding competitor were added in each reaction. U corresponds to the unbound probes and B to the probes bound to MBP-PPR-B.

### PPR-B interferes with the progression of translating ribosomes along the *orf138* mRNA

The location of PPR-B binding site within *orf138* coding sequence led us to analyse in details the distribution of ribosome footprints along the *orf138* coding sequence in the presence and in the absence of PPR-B. Firstly, RNA-Seq read profiles along the di-cistronic *orf138* transcript did not reveal any major different between the CMS and B1 lines (Fig. 6*A*), except a slight decrease of read coverage for both *orf138* and *atp8* in the restored line as shown in Fig. S3. In contrast, Ribo-Seq read distributions confirmed the strong decrease of ribosome coverage on the *orf138* open reading frame and a lack of major impact of PPR-B on *atp8* translation. The position of the PPR-B binding site within *orf138* suggested that it may block translation elongation along the transcript. To see whether this might be possible, we calculated the average number of ribosomes along the *orf138* transcript before and after the PPR-B binding site in both B1 and CMS lines. These data were normalized by both gene-fragment length and abundance, as estimated from RNA-Seq read coverage. A moderate reduction of about 25% in ribosome coverage could be detected in the B1 line before PPR-B binding site (Fig. 6*B*), which is thus much less pronounced than the global translational decrease measured for *orf138* in B1 (Fig. 3*A*). In contrast, the shortage in ribosome density along the *orf138* segment located downstream of PPR-B binding site was found to be much more dramatic (Fig. 6*B*) and in the same magnitude as the observed translation reduction of the *orf138* gene in B1 line (Fig. 3*A*). This observation strongly suggested a normal translational initiation on *orf138* mRNAs in the presence of PPR-B, but rather an incapacity of elongating to cross the PPR-B binding site when the fertility restorer protein is present.

**Figure 6.**
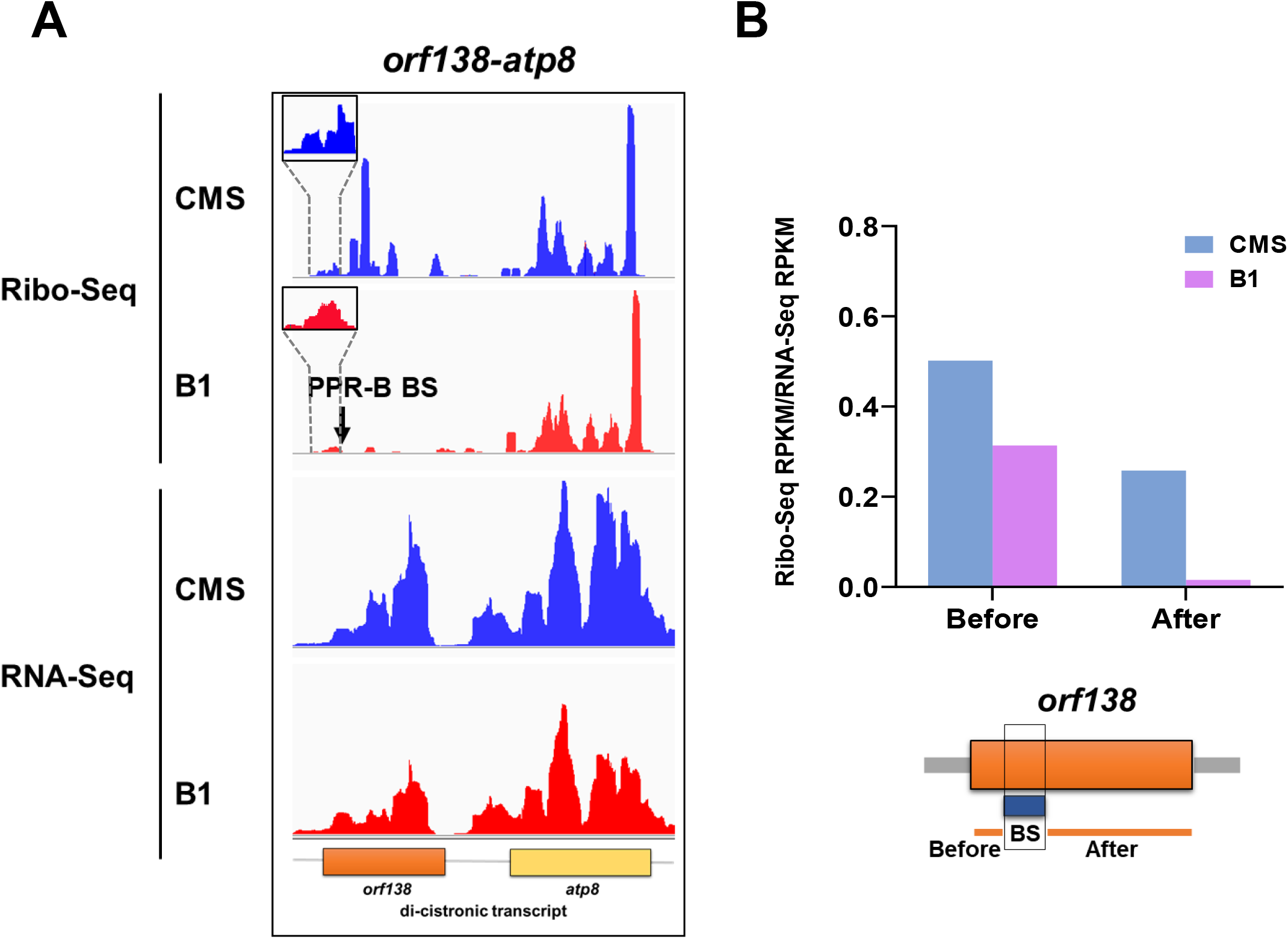
PPR-B-mediated decrease in ribosome footprint along *orf138* is much more pronounced downstream than upstream of PPR-B binding site. (A) Screen captures from the Integrated Genome Viewer software showing the distribution of RNA-Seq reads and ribosome footprints (Ribo-Seq) along the *orf138* locus in B1 and CMS lines. The distributions were normalized to the number of reads mapping to the mitochondrial genome. Zoomed-in views of the ribosome footprint distributions upstream of the PPR-B binding site are shown for both genotypes in upper left corners. (B) Normalized ribosome footprint densities calculated upstream and downstream of PPR-B binding site in the *orf138* transcript. Shown data are means of two biological repeats.

## Discussion

### The Ogura fertility restorer protein is an mRNA-specific translation elongation inhibitor

Besides being essential for protein production, mRNA translation is an important control step in gene expression and all phases of translation constitute potential control checkpoints. It has been shown that the initiation step, which consists in the loading of the small ribosomal subunit (SSU) and the charged initiator tRNA on the start codon prior to full ribosome assembly represents the major checkpoint of translational control (28–31). In eubacteria, such regulation mostly operates by outcompeting the binding of the initiation complex to the mRNA 5’ translation initiation region, most particularly to the Shine-Dalgarno sequence. This can be achieved by multiple ways like the association of regulatory proteins, small antisense RNAs (sRNAs) or metabolites, or *via* the action of signals like temperature or pH (1, 2). In eukaryotes, the regulation of translation initiation occurs mostly *via* the binding of partially-complementary antisense microRNAs (miRNAs) to mRNA 5’ or 3’ UTRs (4). Our analysis reveals that the translation control used to silence the CMS-causing *orf138* transcript of the Ogura system operates at a different level than the initiation step. Translational inhibition has long suspected to be the molecular mechanism associated with fertility restoration in this CMS system since the decrease in Orf138 protein accumulation is not accompanied by any impact on the *orf138* mRNA accumulation, whilst PPR-B was found to associate with *orf138* transcript *in vivo* (22). The *in organello* synthesis, polysome sedimentation and Ribo-Seq analyses presented in this study are perfectly concordant and unambiguously support that PPR-B-mediated *orf138* silencing effectively involves a specific inhibition of *orf138* translation (Figs. 1, 2 and 3). Moreover, the location of PPR-B binding sites within the *orf138* coding sequence (Figs. 4 and 5) and the unchanged ribosome density upstream of this site when PPR-B is present (Fig. 6) strongly support that PPR-B translational control occurs by impeding translation elongation along the *orf138* mRNA and not by affecting the initiation step. PPR-B appears thus to act by blocking ribosome translocation during translation elongation along *orf138* transcript, most likely by steric hindrance. Such translational control was not previously described to take place in plant organelles to control the expression of either essential organellar genes or CMS-associated orfs (11, 32). The mode action of PPR-B is however reminiscent of the way organellar helical repeat proteins, including PPR proteins, act to stabilize mitochondrial and plastid RNAs by impeding the progression of exoribonucleases along mRNAs from their 5’ or 3’ extremities (33–37). PPR protein thus have an inherent capacity to act as roadblocks to impede the progression of RNA processing enzymes along transcripts, which in the case of ribosomes implies to counteract the strong helicase activity of elongating ribosomes (38). The binding strength of PPR-B to its target site seems important to efficiently block translation elongation along the *orf138* mRNAs *in vivo* as we previously showed that all PPR-B repeats are indispensible for complete fertility restoration in rapeseed (39). Similarly, a non-restoring allele of PPR-B comprising only 4 amino acid substitutions was also found to be incompetent for fertility restoration in radish (21). Other known examples involving impairment of translation elongation are scarce. Codon usage, Shine and Dalgarno-like sequences or mRNA secondary structures have been shown to influence translation elongation speed or rate along bacterial orfs but do not lead to a complete block of translation elongation (1). In plants, miRNAs have also been found to partially work as ribosome blocker, although the biological impact of such translational repression remains unclear (4, 40, 41). The analysis of PPR-B activity reveals thus a novel way of translational control which leads to an arrest of the translation elongation for a mitochondrial transcript. The recent deciphering of the PPR recognition code (25, 42) allows the recoding of PPR proteins to bind any sequence of interest (43–45). It will therefore be interesting to see if PPR-B translation inhibitory activity can be recreated using synthetic or recoded PPRs, and then use such PPRs to investigate whether the blockade of translational elongation requires a binding in the proximal region of *orf138* mRNA or not.

**Figure 7.**
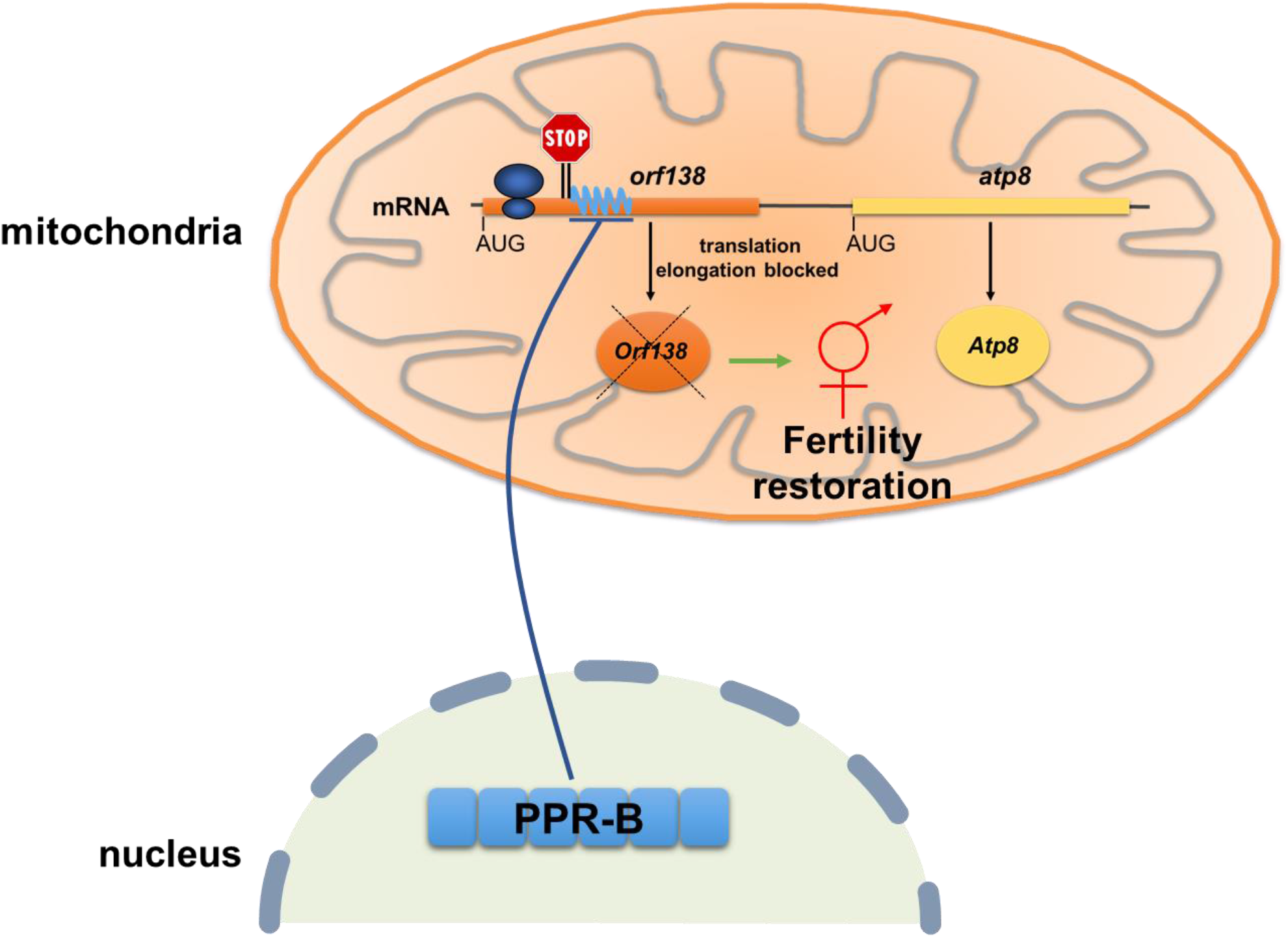
Model of PPR-B mode of action to restore fertility in the Ogura CMS system. Drawing illustrating the molecular mode of action of the Ogura fertility restorer (PPR-B) which, after transfer into mitochondria, specifically inhibits translation elongation along the *orf138* mRNA, thereby inhibiting the production of the mitochondrial protein Orf138 and restoring male fertility.

### The Ogura fertility restorer protein is the first fertility restorer shown to act as a translation elongation blocker

Restorer of fertility genes have been cloned from various crop species and most of them were found to encode PPR proteins (8, 46). Large-scale phylogenetic analyses have shown that identified Rf-PPR genes are evolutionary related and have evolved from distinct subgroup of PPR genes within the PPR family in angiosperms called Rf-like or RFL (47). However, molecular analyses have revealed that Rf-PPRs reduce the accumulation of their cognate CMS-inducing mitochondrial RNAs or proteins through different mechanisms. Anyhow, the vast majority of fertility restorer proteins like RF1A, RF1B or RF6 in rice (12, 48), RFN and RFP in rapeseed (49–51) or RF1 and RF3 in wheat (52) induce specific cleavage within the coding sequence of their cognate CMS-inducing mitochondrial transcripts. Such Rf-PPRs, and several other RFL proteins involved in the processing of conserved mitochondrial transcripts (53–57), generally bind 20-100 bases upstream of the cleavage sites and induce subsequent endonucleolytic processing through a still unclear mechanism (58). The recruitment of an unidentified endonuclease and a potential influence of RNA secondary structures or sequence close to the processing sites have been proposed to explain why cleavage does not occur always at the same distance from PPR binding sites (45). In most cases, the RNA cleavage induced by Rf-PPRs result in a significant decrease in the accumulation of CMS transcripts, leading to a reduction in the production of corresponding CMS proteins and hence to fertility restoration. Therefore, the translational suppression activity that we reveal here for the Ogura fertility restorer is the very first example described for a PPR protein and thus for a PPR-Rf. It has been suggested the rice RF1A may negatively impact the translation of its cognate CMS mRNA, *orf79.* However, the observed translational inhibitory effect is not directly imputable to RF1A but is secondary to an RF1A-induced RNA cleavage liberating a non-translatable monocistronic form of the *orf79* transcript (59). Similarly, the petunia Rf-PPR592 protein has been suggested to impact the translation of the CMS-associated mRNA *pcf*, although changes in *pcf* 5’ processing were also detected in restored plants (60). RNA co-immunoprecipitation assays with Rf-PPR592 showed greatest enrichment in a region of the *pcf* 5’ leader overlapping with the processing site that is altered in restored plants (61). These results do not allow to firmly conclude on the role of Rf-PPR592 in fertility restoration but its preferential association with a region of *pcf* 5’ UTR favours a role in *pcf* transcript 5’-end processing (62). It also remains possible that this processing prevents proper translation of *pcf* mRNA, implying an indirect role of Rf-PPR592 in *pcf* translation. The reason why PPR-B binding does not induce any cleavage within the *orf138* transcript is currently unclear. PPR-B sequence is highly similar to that of RFL proteins known to induce endonucleolytic cleavage in Arabidopsis (47, 53). Minor sequence differences between PPR-B and these RFLs or the involvement of specific *cis*-elements or RNA secondary structures downstream of the binding sites may be responsible for their difference in activity. The molecular mode of action of PPR-B demonstrates that CMS genes could be silenced by simply targeting a PPR protein in their coding sequence to inhibit their translation, regardless of the presence of sequences or structural elements favourable for RNA cleavage. Our observations should thus facilitate the production of synthetic fertility restorers for CMS systems in which no efficient restorers have been identified up to now.

## SI Materials and Methods

### Plant material

Rapeseed (*Brassica napus*) plants were grown in the greenhouse at 20-25°C, under a 16 h light/8 h dark cycle. The B1, CMS (Pactol Sterile) lines used in this study were previously described in (22). For polysome sedimentation experiments, a B1 plant and its non-transgenic parental line (Pactol Fertile) were used to pollinate the male-sterile 18S cybrid (24) and the analysis was developed on generated F1 hybrids.

### Preparation of mitochondria

Mitochondria were isolated from rapeseed floral buds as previously described in (22). Briefly, flower buds were blended in grinding buffer containing 300 mM sucrose, 25 mM tetrasodium pyrophosphate, 10 mM KH2PO4, 2 mM EDTA, 0.8% (w/v) polyvinylpyrrolidone-40, 0.3% [w/v] BSA, and 20 mM ascorbate, pH 7.5. The homogenate was filtered through three layers of Miracloth membrane (Calbiochem), followed by three successive low-speed centrifugations at 2000, 2600, and 3000 *g* for 10 min at 4°C. The supernatant was recovered and centrifuged at 23,400 *g* for 20 min to pellet a fraction enriched in mitochondria. The pellet was then resuspended in washing buffer (0.3 M sucrose and 10 mM HEPES-KOH, pH 7.5), loaded on a 14-25-50% (v/v) Percoll step gradient diluted in washing buffer supplemented with 0.2% BSA and centrifuged for 20 min at 24,000 *g*. Mitochondria were collected from the 50/25% interface, diluted at least 10 times in washing buffer, and pelleted at 23,400 *g* for 20 min for subsequent use.

### Polysome association analysis

Polysome analyses were performed on flower bud extracts as described previously (63). After ultra-centrifugation, ten 900 μL fractions were manually collected from the top of the gradients. Total RNA was prepared from each of them and analysed by RNA-gel blot analysis using probes recognizing *orf138*, *atp9* and *atp1* transcripts. The primers used to prepare these different DNA probes are indicated in Supplementary Table S1.

### Immunodetection of proteins

Mitochondrial proteins were extracted in 30 mM HEPES-KOH (pH 7.7), 10 mM Mg(OAc)2, 150 mM KOAc, 10% glycerol and 0.5% (w/v) CHAPS. Protein concentrations were measured using the Bradford reagent (Bio-Rad) and separated by SDS-PAGE (polyacrylamide gel electrophoresis). After electrophoresis, gels were transferred onto PVDF membranes (Bio-Rad) under semidry conditions. Membranes were hybridized with antibodies using dilutions indicated in Table S2.

### Ribo-Seq and RNA-Seq library preparation and sequencing

Mitoribosome footprints were prepared from rapeseed flower buds as previously described (64). Ribosome footprints were depleted from ribosomal RNA with the Ribo-Zero Plant kit (Illumina) following manufacturer’s recommendations. Ribo-Seq libraries were prepared using the TruSeq Small RNA library preparation kit (Illumina). For RNA-Seq, total RNAs were extracted from 1/10th of the lysates used for Ribo-Seq analysis and treated with the Ribo-Zero rRNA Removal Kit (Illumina). One hundred ng of the rRNA-depleted RNAs were used for library construction using the NEXTflex Rapid Directional qRNA-Seq Kit (Bioo Scientific) according the manufacturer’s instructions. Next generation sequencing was performed on a HiSeq 2500 instrument (Illumina) for Ribo-Seq libraries (single end, 50 nt) or NextSeq 500 sequencer (Illumina) for RNA-Seq libraries (single end, 75 nt).

### Bioinformatic analyses

Ribo-Seq and RNA-Seq sequencing data were processed and mapped as previously described in (64). Ribo-Seq and RNA-Seq RPKMs were calculated based on reads mapping to mitochondrial and nuclear coding sequences following a procedure detailed in (65) and translation efficiencies were calculated as ratios of ribosome footprint RPKMs to RNA-seq RPKMs. *B. napus* mitochondrial (GenBank AP006444.1) and nuclear (GenBank GCF_000686985.2) genome sequences were used for read mapping. To permit read mapping along the *orf138* locus as well, its sequence (18), comprising both *orf138* and *atp8* open reading frames, was manually added to that of the *B. napus* mitochondrial genome. The coding sequence of *orf138* was however truncated by 138 nucleotides on its 3’ end to remove the three perfect repeats found at the end of the gene (*SI Appendix*, Fig. S4). Consequently, the *B. napus* endogenous *atp8* sequence was removed from the generated mitochondrial genome to avoid duplicated *atp8* gene copies. Similarly, the pseudogenized and nonconserved copy of *cox2 (cox2-2* (66, 67)) was removed from the AP006444.1 mitochondrial genome to avoid a lack of read mapping along the *cox2* gene. An adapted .gff annotation file taking into account these different modifications was created for mapped read counting.

### Gel mobility shift assays

A truncated version of PPR-B protein deprived of its mitochondrial transit peptide but comprising all PPR repeats was expressed in fusion with a N-terminal maltose binding protein (MBP) protein in *E. coli* Rosetta DE3 cells (Novagen). The corresponding *PPR-B* DNA fragment was PCR amplified using the GWPPRB-5 and GWPPRB-3 primers (Table S1). The obtained amplification was subcloned into pDONR207 by Gateway™ BP reaction (Invitrogen) and subsequently transferred into pMAL-TEV (37) by Gateway™ LR reaction (Invitrogen). Following expression at 37°C for 3 h, obtained bacterial pellets were lysed in 50 mM HEPES-KOH (pH 7.8), 150 mM NaCl, 1% glycerol, 0.01% CHAPS and 5 mM β-mercaptoethanol using the One-Shot cell disruption system (Constant Systems). After centrifugation at 20,000 *g* for 15 min, the MBP-PPR-B fusion protein was purified from the supernatant on amylose beads (Biolabs) following manufacturer’s recommendations. Protein purity was verified on SDS-PAGE before proceeding to Gel mobility shift (GMS) assays. Gel mobility shift assays were performed as previously described (37) using *in vitro*-transcribed radiolabeled RNA probes. Binding reactions were performed in 25 mM HEPES-KOH (pH 7.5), 150 mM NaCl, 0.1 mg/ml BSA, 10% glycerol, 0.5 mg/ml heparin, 2 mM DTT, containing 200 pM of radiolabeled RNA probes and purified MBP-PPR-B at 200 nM, 400 nM, or 800 nM. Binding reactions were incubated for 45 min at 25°C and resolved on a non-denaturing 5% polyacrylamide gel at 150 volts in 1 × THE (66 mM HEPES, 34 mM Tris (pH 7.7), 0.1 mM EDTA). The data were visualized with a phosphorimager (FLA-9500 Fujifilm).

### *In organello* protein synthesis

*In organello* protein synthesis experiments were done as previously described in (24).

## ACKNOWLEDGMENTS

This work was supported by the Agence Nationale de la Recherche (ANR) MITRA Grant ANR-16-CE11-0024-01 (to H.M.) and the China Scholarship Council (to C.W.). The Institut Jean-Pierre Bourgin’s (IJPB’s) benefits from the support of Saclay Plant Sciences Grant ANR-17-EUR-0007. This work has benefited from the support of IJPB’s Plant Observatory technological platforms.

## Author contributions

H.M. designed research. C.W., L.L., N.A. and M.Q. performed research. C.W. and H.M. analysed the data. C.W. and H.M. wrote the paper.

---

**Supplemental Figure 1.**
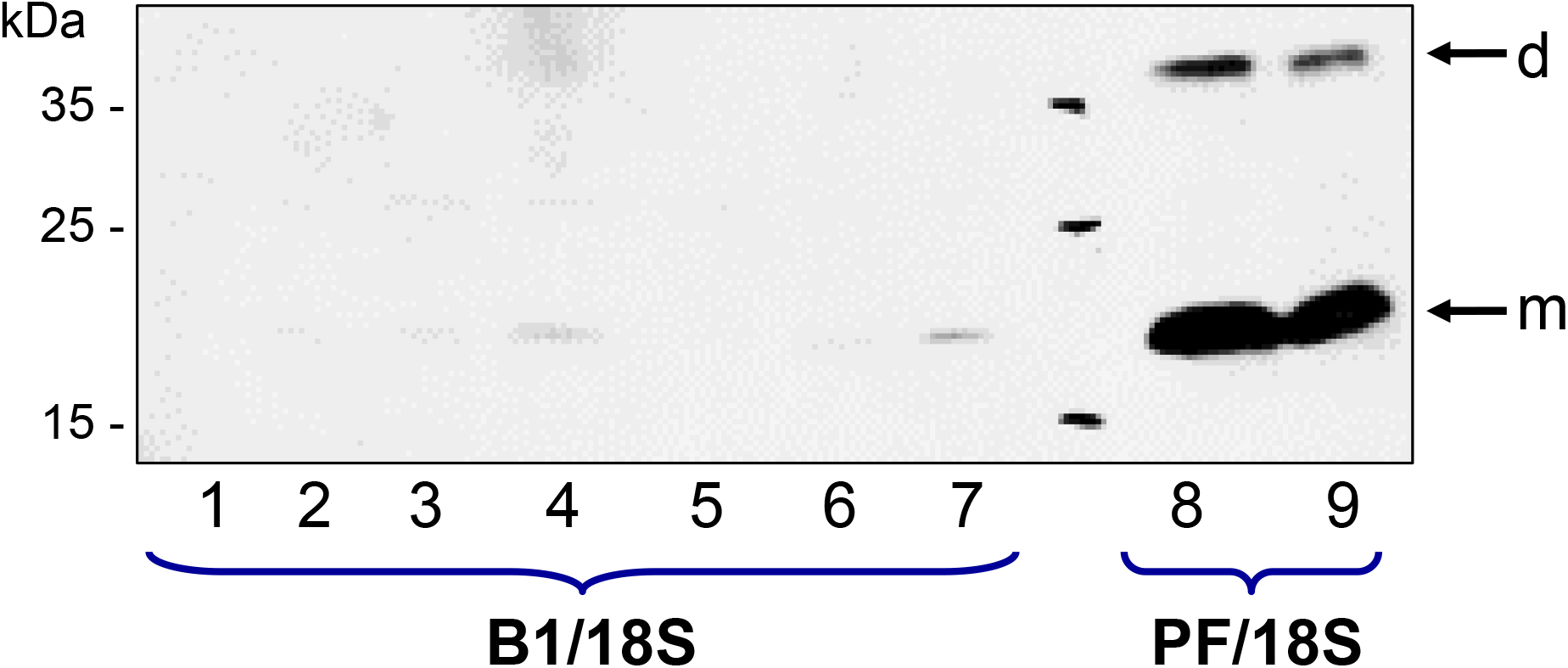
Immunoblot analysis of mitochondrial proteins exacted from B1/18S and Pactol Fertile/18S hybrid plants to evaluate the accumulation level of Orf138 in these plants. m: monomeric Orf138, d: dimeric Orf138.

**Supplemental Figure 2.**
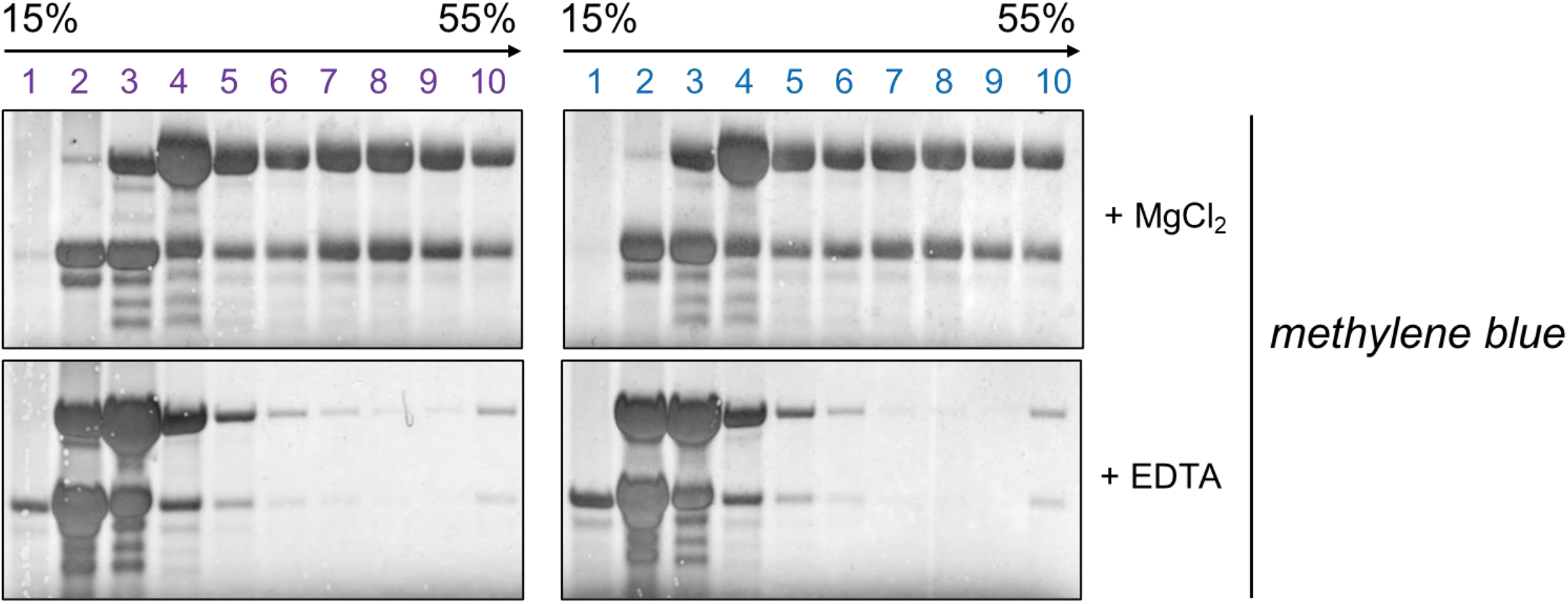
RNA gel blot stained with methylene blue revealing the RNA content of fractions recovered from a representative polysome sedimentation experiment. Flower bud RNA extracts were fractionated in 15-55% continuous sucrose density gradients by ultracentrifugation under conditions either maintaining (MgCl_2_) or disrupting (EDTA) polysome integrity. The obtained profiles indicate that polysomal RNAs migrate towards the centre and the bottom of the gradients, whereas free mRNAs accumulate in the upper fractions.

**Supplemental Figure 3.**
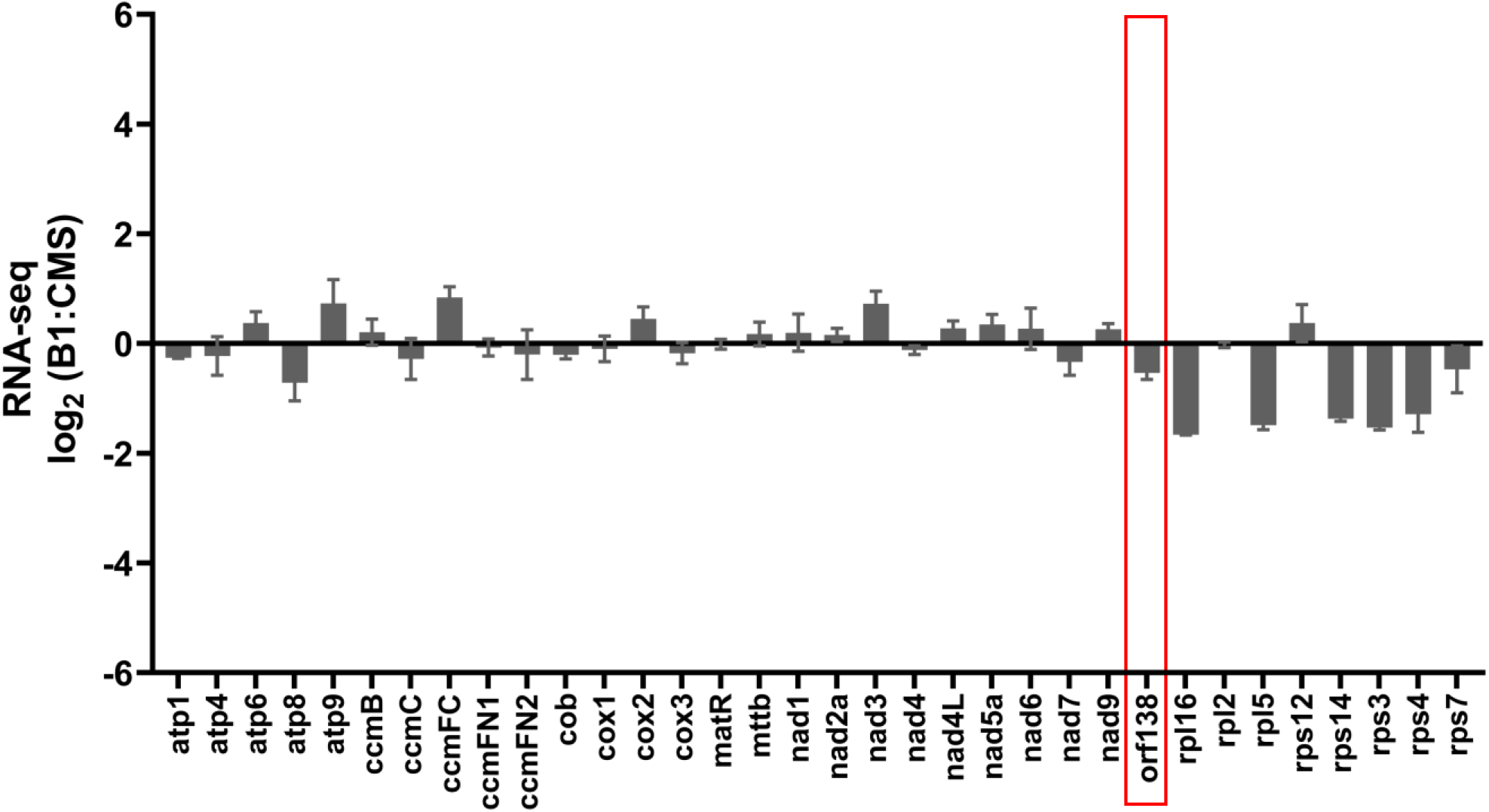
Genome-wide view of mitochondrial mRNA abundance in B1 versus CMS plants, as estimated by RNA-Seq analysis. The data represent reads per kilobase per million reads mapping to nuclear genome coding sequences (RPKM). Shown values are B1 to CMS ratios for each mitochondrial gene and are means ± SD from three biological replicates.

**Supplemental Figure 4.**
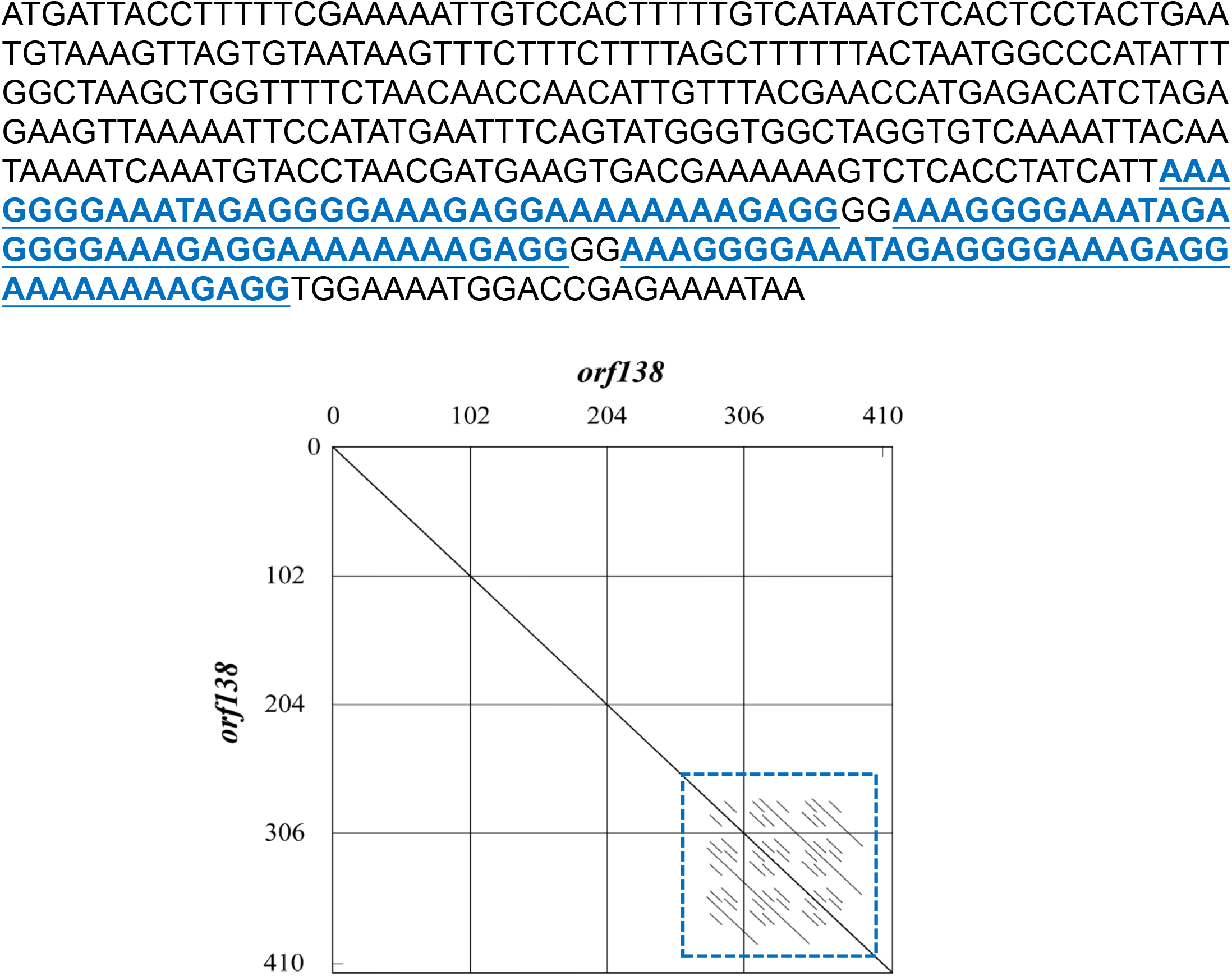
Sequence of the *orf138* gene showing the three repeated sequences present at the end of the gene. The repeated sequence is marked in blue and is underlined. The box plot below shows that no other repeated sequence is found in the *orf138* gene.

**Supplemental Table S1.**
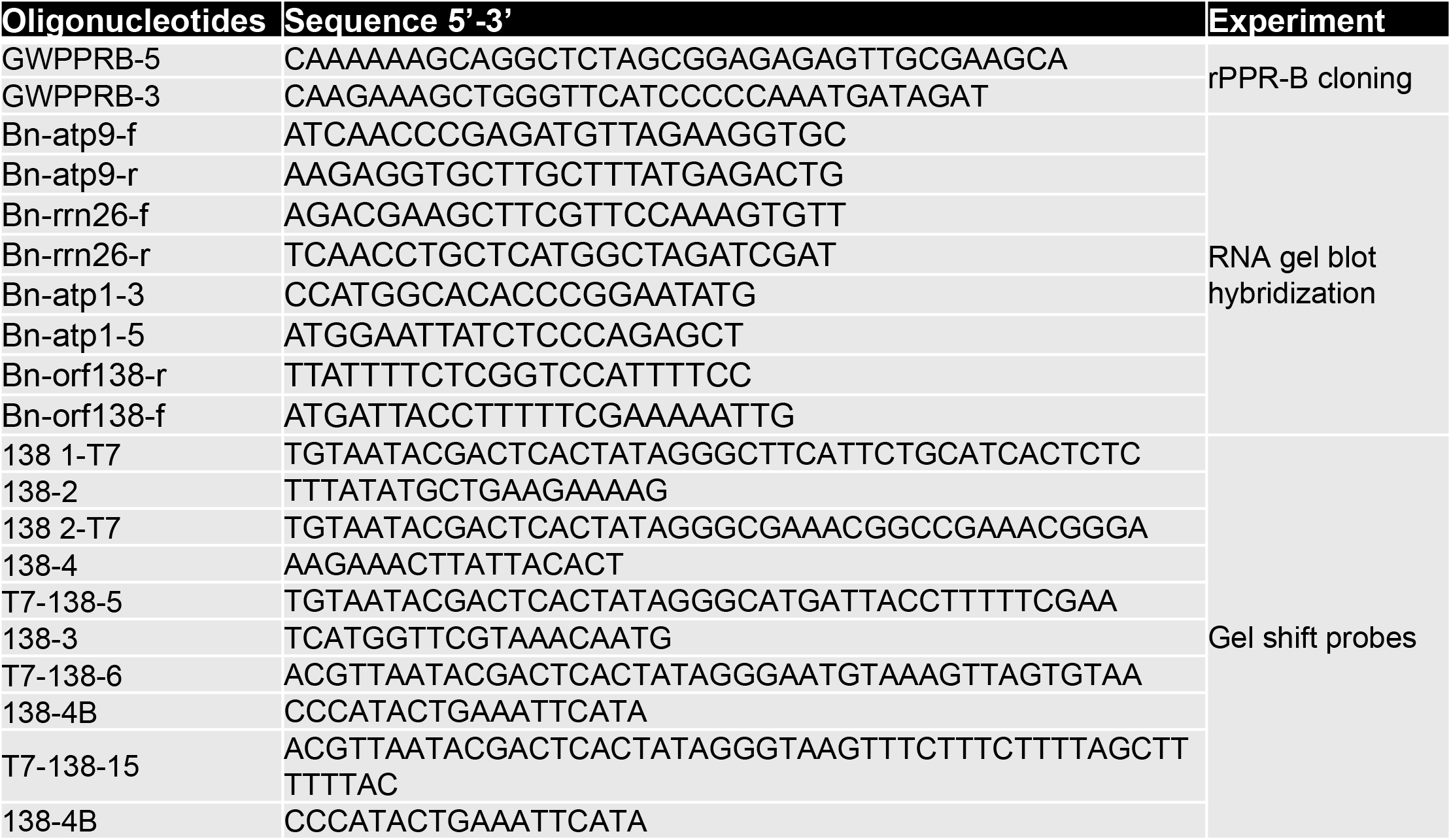
Oligonucleotides used in this study.

**Supplemental Table S2.** List of antibodies used in this study.

| Antibody | Full Name of detected proteins | Host | Dilution | Source |
| --- | --- | --- | --- | --- |
| Orf138 | Open reading frame 138 | Rabbit | 1 : 2,000 | (1) |
| PPRB | pentatricopeptide repeat protein B | Rabbit | 1 : 1,000 | (2) |
| Nad6 | NADH-ubiquinone oxidoreductase chain 6 | Rabbit | 1 : 1,000 | Agrisera (AS15 2926) |
| Nad7 | NADH-ubiquinone oxidoreductase chain 7 | Rabbit | 1 : 2,000 | (3) |
| Nad9 | NADH-ubiquinone oxidoreductase chain 9 | Rabbit | 1 : 5,000 | (4) |
| RISP | Rieske iron-sulfur protein | Rabbit | 1 : 5,000 | (5) |
| Cox2 | Cytochrome oxidase subunit II | Rabbit | 1 : 1,000 | Agrisera (AS04 053A) |
| Rpl16 | Mitochondrial ribosomal large subunit protein L16 | Rabbit | 1 : 1,000 | Agrisera (AS15 3069) |
| Porin | The channel-forming protein | Mouse | 1 : 500 | (6) |

## References

1. Chiaruttini C, Guillier M (2020) On the role of mRNA secondary structure in bacterial translation. WIREs RNA 11(3):7007.

2. Babitzke P, Lai Y-J, Renda AJ, Romeo T (2019) Posttranscription Initiation Control of Gene Expression Mediated by Bacterial RNA-Binding Proteins. Annu Rev Microbiol 73(1):43–67.

3. Neelagandan N, Lamberti I, Carvalho HJF, Gobet C, Naef F (2020) What determines eukaryotic translation elongation: recent molecular and quantitative analyses of protein synthesis. Open Biol 10(12):200292.

4. Iwakawa H-O, Tomari Y (2015) The Functions of MicroRNAs:mRNA Decay andTranslational Repression. Trends in Cell Biology 25(11):651–665.

5. Hammani K, Giegé P (2014) RNA metabolism in plant mitochondria. Trends in Plant Science 19(6):380–389.

6. Dennerlein S, Wang C, Rehling P (2017) Plasticity of MitochondrialTranslation. Trends in Cell Biology 27(10):712–721.

7. Barkan A (2011) Expression of plastid genes: organelle-specific elaborations on a prokaryotic scaffold. Plant Physiology 155(4):1520–1532.

8. Chen L, Liu Y-G (2014) Male Sterility and Fertility Restoration in Crops. Annu Rev Plant Biol 65(1):579–606.

9. Gaborieau L, Brown GG, Mireau H (2016) The Propensity of Pentatricopeptide Repeat Genes to Evolve into Restorers of Cytoplasmic Male Sterility. Front Plant Sci 7(e1002910):271.

10. Lurin C, et al. (2004) Genome-wide analysis of Arabidopsis pentatricopeptide repeat proteins reveals their essential role in organelle biogenesis. Plant Cell 16(8):2089–2103.

11. Barkan A, Small I (2014) Pentatricopeptide Repeat Proteins in Plants. Annu Rev Plant Biol 65(1):415–442.

12. Wang Z, et al. (2006) Cytoplasmic male sterility of rice with boro II cytoplasm is caused by a cytotoxic peptide and is restored by two related PPR motif genes via distinct modes of mRNA silencing. Plant Cell 18(3):676–687.

13. Bentolila S, Alfonso AA, Hanson MR (2002) A pentatricopeptide repeat-containing gene restores fertility to cytoplasmic male-sterile plants. Proc Natl Acad Sci USA 99(16):10887–10892.

14. Tang H, et al. (2014) The Rice Restorer Rf4 for Wild-Abortive Cytoplasmic Male Sterility Encodes a Mitochondrial-Localized PPR Protein that Functions in Reduction of WA352 Transcripts. Molecular Plant 7(9):1497–1500.

15. Igarashi K, Kazama T, Toriyama K (2016) A Gene Encoding Pentatricopeptide Repeat Protein Partially Restores Fertility in RT98-Type Cytoplasmic Male-Sterile Rice. Plant Cell Physiol 57(10):2187–2193

16. Bellaoui M, Grelon M, Pelletier G, Budar F (1999) The restorer Rfo gene acts post-translationally on the stability of the ORF138 Ogura CMS-associated protein in reproductive tissues of rapeseed cybrids. Plant Mol Biol 40(5):893–902.

17. Bonhomme S, Budar F, Férault M, Pelletier G (1991) A 2.5 kb NcoI fragment of Ogura radish mitochondrial DNA is correlated with cytoplasmic male-sterility in Brassica cybrids. Curr Genet 19:121–127.

18. Bonhomme S, et al. (1992) Sequence and transcript analysis of the Nco2.5 Ogura-specific fragment correlated with cytoplasmic male sterility in Brassica cybrids. Mol Gen Genet 235(2-3):340–348.

19. Desloire S, et al. (2003) Identification of the fertility restoration locus, Rfo, in radish, as a member of the pentatricopeptide-repeat protein family. EMBO Rep 4(6):588–594.

20. Brown GG, et al. (2003) The radish Rfo restorer gene of Ogura cytoplasmic male sterility encodes a protein with multiple pentatricopeptide repeats. The Plant Journal 35(2):262–272.

21. Koizuka N, et al. (2003) Genetic characterization of a pentatricopeptide repeat protein gene, orf687, that restores fertility in the cytoplasmic male-sterile Kosena radish. Plant J 34(4):407–415.

22. Uyttewaal M, et al. (2008) Characterization of Raphanus sativus Pentatricopeptide Repeat Proteins Encoded by the Fertility Restorer Locus for Ogura Cytoplasmic Male Sterility. Plant Cell 20(12):3331–3345.

23. Hanson M, Bentolila S (2004) Interactions of mitochondrial and nuclear genes that affect male gametophyte development. Plant Cell 16 Suppl:S154–69.

24. Grelon M, Budar F, Bonhomme S, Pelletier G (1994) Ogura cytoplasmic male-sterility (CMS)-associated orf138 is translated into a mitochondrial membrane polypeptide in male-sterile Brassica cybrids. Mol Gen Genet 243(5):540–547.

25. Barkan A, et al. (2012) A combinatorial amino Acid code for RNA recognition by pentatricopeptide repeat proteins. PLoS Genet 8(8):e1002910.

26. Takenaka M, Zehrmann A, Brennicke A, Graichen K (2013) Improved Computational Target Site Prediction for Pentatricopeptide Repeat RNA Editing Factors. PLoS ONE 8(6):e65343.

27. Yagi Y, Hayashi S, Kobayashi K, Hirayama T, Nakamura T (2013) Elucidation of the RNA Recognition Code for Pentatricopeptide Repeat Proteins Involved in Organelle RNA Editing in Plants. PLoS ONE 8(3):e57286.

28. Merrick WC, Pavitt GD (2018) Protein Synthesis Initiation in Eukaryotic Cells. Cold Spring Harbor Perspectives in Biology 10(12). doi:10.1101/cshperspect.a033092.

29. Gualerzi CO, Pon CL (2015) Initiation of mRNA translation in bacteria: structural and dynamic aspects. Cell Mol Life Sci 72(22):4341–4367.

30. Shah P, Ding Y, Niemczyk M, Kudla G, Plotkin JB (2013) Rate-Limiting Steps in Yeast Protein Translation. Cell 153(7):1589–1601.

31. Gebauer F, Hentze MW (2004) Molecular mechanisms of translational control. Nature Reviews Molecular Cell Biology 5(10):827–835.

32. Kim Y-J, Zhang D (2018) Molecular Control of Male Fertility for Crop Hybrid Breeding. Trends in Plant Science 23(1):53–65.

33. Prikryl J, Rojas M, Schuster G, Barkan A (2011) Mechanism of RNA stabilization and translational activation by a pentatricopeptide repeat protein. Proc Natl Acad Sci USA 108(1):415–420.

34. Pfalz J, Bayraktar OA, Prikryl J, Barkan A (2009) Site-specific binding of a PPR protein defines and stabilizes 5” and 3” mRNA termini in chloroplasts. The EMBO Journal 28(14):2042–5211.

35. Hammani KK, Cook WBW, Barkan AA (2012) RNA binding and RNA remodeling activities of the half-a-tetratricopeptide (HAT) protein HCF107 underlie its effects on gene expression. Proc Natl Acad Sci USA 109(15):5651–5656.

36. Haïli N, et al. (2013) The pentatricopeptide repeat MTSF1 protein stabilizes the nad4 mRNA in Arabidopsis mitochondria. Nucleic Acids Res 41(13):6650–6663.

37. Wang C, et al. (2017) The pentatricopeptide repeat protein MTSF2 stabilizes a nad1 precursor transcript and defines the 3’ end of its 5’-half intron. Nucleic Acids Res 45(10):6119–6134.

38. Takyar S, Hickerson RP, Noller HF (2005) mRNA helicase activity of the ribosome. Cell 120(1):49–58.

39. Qin X, et al. (2014) In vivo functional analysis of a nuclear restorer PPR protein. BMC Plant Biol 14:313.

40. Iwakawa H-O, Tomari Y (2013) Molecular insights into microRNA-mediated translational repression in plants. Mol Cell 52(4):591–601.

41. Chung BY-W, Deery MJ, Groen AJ, Howard J, Baulcombe DC (2017) Endogenous miRNA in the green alga Chlamydomonas regulates gene expression through CDS-targeting. Nature Plants: 3(10):787–794.

42. Shen C, et al. (2016) Structural basis for specific single-stranded RNA recognition by designer pentatricopeptide repeat proteins. Nature Communications 7:11285.

43. Yu Q, Barkan A, Maliga P (2019) Engineered RNA-binding protein for transgene activation in non-green plastids. Nature Plants: 5(5):486–490.

44. Rojas M, Yu Q, Williams-Carrier R, Maliga P, Barkan A (2019) Engineered PPR proteins as inducible switches to activate the expression of chloroplast transgenes. Nature Plants: 5(5):505–511.

45. Francs-Small des CC, Sanglard LVP, Small I (2018) Targeted cleavage of nad6 mRNA induced by a modified pentatricopeptide repeat protein in plant mitochondria. Communications Biology: 1:166.

46. Dahan J, Mireau H (2013) The Rf and Rf-like PPR in higher plants, a fast-evolving subclass of PPR genes. RNA Biol 10(9):1469–1476.

47. Fujii S, Bond CS, Small ID (2011) Selection patterns on restorer-like genes reveal a conflict between nuclear and mitochondrial genomes throughout angiosperm evolution. Proc Natl Acad Sci USA 108(4):1723–1728.

48. Huang W, et al. (2015) Pentatricopeptide-repeat family protein RF6 functions with hexokinase 6 to rescue rice cytoplasmic male sterility. Proceedings of the National Academy of Sciences. doi:10.1073/pnas.1511748112.

49. Singh M, et al. (1996) Nuclear genes associated with a single Brassica CMS restorer locus influence transcripts of three different mitochondrial gene regions. Genetics 143(1):505–516.

50. Gaborieau L, Brown GG (2016) Comparative genomic analysis of the compound Brassica napus Rf locus. BMC Genomics 17(1):834.

51. Liu Z, et al. (2016) A Mitochondria-Targeted PPR Protein Restores pol Cytoplasmic Male Sterility by Reducing orf224 Transcript Levels in Oilseed Rape. Molecular Plant 9(7):1082–1084.

52. Melonek J, et al. (2021) The genetic basis of cytoplasmic male sterility and fertility restoration in wheat. Nature Communications: 12(1):1036.

53. Arnal N, Quadrado M, Simon M, Mireau H (2014) A restorer-of-fertility like pentatricopeptide repeat gene directs ribonucleolytic processing within the coding sequence of rps3-rpl16 and orf240a mitochondrial transcripts in Arabidopsis thaliana. Plant J 78(1):134–145.

54. Hölzle A, et al. (2011) A RESTORER OF FERTILITY-like PPR gene is required for 5’-end processing of the nad4 mRNA in mitochondria of Arabidopsis thaliana. Plant J 65(5):737–744.

55. Hauler A, et al. (2013) RNA PROCESSING FACTOR 5 is required for efficient 5’ cleavage at a processing site conserved in RNAs of three different mitochondrial genes in Arabidopsis thaliana. Plant J 74(4):593–604.

56. Jonietz C, Forner J, Hölzle A, Thuss S, Binder S (2010) RNA PROCESSING FACTOR2 is required for 5’ end processing of nad9 and cox3 mRNAs in mitochondria of Arabidopsis thaliana. Plant Cell 22(2):443–453.

57. Jonietz C, Forner J, Hildebrandt T, Binder S (2011) RNA PROCESSING FACTOR3 is crucial for the accumulation of mature ccmC transcripts in mitochondria of Arabidopsis accession Columbia. Plant Physiology 157(3):1430–1439.

58. Binder S, Stoll K, Stoll B (2016) Maturation of 5’ ends of plant mitochondrial RNAs. Physiol Plant 157(3):280–288.

59. Kazama T, Nakamura T, Watanabe M, Sugita M, Toriyama K (2008) Suppression mechanism of mitochondrial ORF79 accumulation by Rf1 protein in BT-type cytoplasmic male sterile rice. Plant J 55(4):619–628.

60. Pruitt KD, Hanson MR (1991) Transcription of the Petunia mitochondrial CMS-associated Pcf locus in male sterile and fertility-restored lines. Mol Gen Genet 227(3):348–355.

61. Gillman JD, Bentolila S, Hanson MR (2007) The petunia restorer of fertility protein is part of a large mitochondrial complex that interacts with transcripts of the CMS-associated locus. Plant J 49(2):217–227.

62. Young EG, Hanson MR (1987) A fused mitochondrial gene associated with cytoplasmic male sterility is developmentally regulated. Cell 50(1):41–49.

63. Barkan A (1993) Nuclear Mutants of Maize with Defects in Chloroplast Polysome Assembly Have Altered Chloroplast RNA Metabolism. Plant Cell 5(4):389–402.

64. Planchard N, et al. (2018) The translational landscape of Arabidopsis mitochondria. Nucleic Acids Res 283:1476.

65. Chotewutmontri P, Alice Barkan (2016) Dynamics of Chloroplast Translation during Chloroplast Differentiation in Maize. PLoS Genet 12(7):e1006106.

66. Handa H (2003) The complete nucleotide sequence and RNA editing content of the mitochondrial genome of rapeseed (Brassica napus L.): comparative analysis of the mitochondrial genomes of rapeseed and Arabidopsis thaliana. Nucleic Acids Res 31(20):5907–5916.

67. Wang J, et al. (2012) Complete sequence of heterogenous-composition mitochondrial genome (Brassica napus) and its exogenous source. BMC Genomics 13(1):1–1.

## References

1. Grelon, M., Budar, F., Bonhomme, S. et al. Ogura cytoplasmic male-sterility (CMS)-associated orf138 is translated into a mitochondrial membrane polypeptide in male-sterile Brassica cybrids. Molec. Gen. Genet. 243, 540–547 (1994).

2. M. Uyttewaal et al., Characterization of Raphanus sativus pentatricopeptide repeat proteins encoded by the fertility restorer locus for Ogura cytoplasmic male sterility. Plant cell. 20, 3331–3345 (2008).

3. B. Pineau, O. Layoune, A. Danon, R. De Paepe, L-galactono-1,4-lactone dehydrogenase is required for the accumulation of plant respiratory complex I. J. Biol. Chem. 283, 32500–32505 (2008).

4. L. Lamattina, D. Gonzalez, J. Gualberto, J. M. Grienenberger, Higher plant mitochondria encode an homologue of the nuclear-encoded 30-kDa subunit of bovine mitochondrial complex I. FEBS J. 217, 831–838 (1993).

5. C. Carrie et al., Conserved and novel functions for Arabidopsis thaliana MIA40 in assembly of proteins in mitochondria and peroxisomes. J. Biol. Chem. 285, 36138–36148 (2010).

6. N. L. Taylor et al., Environmental stresses inhibit and stimulate different protein import pathways in plant mitochondria. FEBS Lett. 547, 125–130 (2003).

